# DUMMEST: A simpler and faster model for frameshift-aware alignment of a DNA sequence to a protein family

**DOI:** 10.64898/2026.10.14.759091

**Authors:** Harvey Shawn Chua, Anish M. S. Shrestha

## Abstract

**Motivation:** We address the problem of aligning DNA sequences containing frameshift errors to protein profile hidden Markov models (pHMMs), a task relevant to annotating protein-coding regions in long-read genomic, metagenomic, and transcriptomic sequence data. Existing approaches either translate sequences across six reading frames, ignoring frameshifts, or rely on slow and unreliable error correction. BATH natively models frameshifts but uses a more complex architecture than conventional pHMMs.

**Results:** We propose a simpler frameshift-aware pHMM based on DUMMER, incorporating modern heuristics, including GPU-accelerated MMseqs2 prefiltering. Evaluations on simulated and real long reads show that our method detects more alignments, with greater average alignment lengths, than non-frameshift-aware tools. Compared with BATH, our method achieves comparable sensitivity and alignment lengths, improved specificity under simulated sequencing artifacts, and is 2–12× faster depending on dataset.

**Availability and Implementation:** **Source code and container image are freely available for download at**: https://github.com/bioinfodlsu/dummest

## Introduction

We consider the following problem:

- **Input:** A reference protein family represented as a profile hidden Markov model (pHMM), and a query DNA sequence likely containing frameshift mutations/errors.
- **Output:** Frameshift-tolerant local alignment(s) between the query and the profile, a score and its statistical significance for each alignment.

Important applications of this problem arise in annotating protein-coding sequences in long-read sequencing data, which are becoming increasingly prevalent across genomics, transcriptomics, and metagenomics. Examples include identifying antibiotic resistance genes and classifying protein domains in metagenomic reads, identifying open reading frames in RNA-seq reads.

The annotation process often involves comparing DNA sequences to reference databases, preferably those containing pHMMs since profile-based alignment is known to be more sensitive than pairwise sequence alignment. However, high error rates that introduce frameshifts in long reads complicate this process. Existing methods generally require error correction or translation before alignment, both of which can reduce sensitivity or introduce additional errors. Therefore we aim to directly align uncorrected, frameshift-error-prone DNA sequences to reference protein family profiles.

**Received:** Date Month Year. **Revised received:** Date Month Year. **Accepted:** Date Month Year

© The Author(s) YEAR. Copyright and licence statements to be updated by the publisher during production.

One of the most widely used pHMM tools is HMMER3 [4], popular due to its high speed and superior sensitivity compared to pairwise alignment tools. Recently, nail [13] introduced an additional speedup over HMMER3 by replacing HMMER3’s prefilters with the more modern MMseqs2 [10] and then computing only the high-scoring portion of the forward-backward matrix based on MMseqs2’s alignments. However, both are designed for aligning protein sequences to protein profiles, and as such lack the ability to align query DNA sequences in a frameshift-aware manner.

BATH [11] builds on HMMER3 by adding frameshift-awareness, albeit with complexity. Conventional protein pHMMs emit 20 symbols per position. In contrast, BATH emits “quasi-codons” (codons containing indels) of lengths 1–5, yielding a total of 4^5^ + 4^4^ + 4^3^ + 4^2^ + 4^1^ = 1364 symbols per position. This 68-fold increase drastically expands the position-specific emission probability tables. For a profile of 500 positions using probabilities stored in the fp32 floating point format, a conventional model requires approximately 40 kB, fitting easily within modern CPU L1 data caches (typically 32–64 kB). BATH, however, requires approximately 2.6 MiB which might compete with other memory accesses and trigger cache misses.

Fast frameshift-aware pairwise alignment tools do exist, including LAST [15] and DIAMOND [3]. However, they cannot exploit position-specific probabilities and generally achieve lower sensitivity than profile-based methods, particularly for distant homologs.

Recently, Frith introduced DUMMER as a simpler alternative to HMMER [8]. DUMMER addresses several limitations of the Plan7 architecture used by HMMER3, which is also shared by nail and BATH. These include a scoring scheme that is biased toward longer alignments and inaccuracies in E-value estimation identified in HMMER3’s own studies [4]. By eliminating alignment length bias, DUMMER is able to leverage the universal significance statistics established by Yu and Hwa [8; 17]. Empirical benchmarks demonstrate that DUMMER achieves sequence search sensitivity and specificity comparable to HMMER3.

Here we describe a frameshift-aware pHMM that draws on the length-bias-free alignment idea of DUMMER [8] and the probability model structure for DNA-protein pairwise alignment used by LAST [15]. A major difference from BATH is that: rather than adding a huge number of states that handle a variety of frameshifted codons (“quasi-codons”), we add nucleotide-emitting indel states to DUMMER, yielding a simpler and faster model. The hypothesis is that combinations of nucleotides and codons can approximate most indels, including those inducing frameshifts. Its simpler architecture is conjectured to follow the universal E-value statistics established by Yu and Hwa [17]. To further reduce runtime, our method employs MMseq2 for multi-stage heuristic prefiltering.

## Methods

### Our profile HMM Model

Our proposed pHMM represents protein family information as repetitive blocks of match, insertion, and deletion states. The full model specification is provided in Supplementary Material Section S1. Briefly, each block *i* contains a match state *M*^*i*^, three insertion states 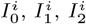, and three deletion states 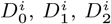, (see Fig. 1). The match state *M*^*i*^ emits a codon whose probability is determined by the amino acid at that profile position. The insertion states model additional nucleotides relative to the profile: 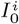 emits a full codon, while 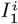 and 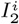 emit one and two nucleotides, respectively, corresponding to frameshifting insertion events. The deletion states allow skipping profile positions: 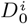 skips a full codon, while 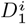 and 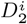 emit two and one nucleotides respectively, accounting for DNA deletions that are not multiples of three. Example alignments through these states are illustrated in Fig. 2. Auxiliary states (shown in gray in Fig. S1) connect each block to the unaligned flanks, allowing alignments to start and end at any block.

**Figure 1.**
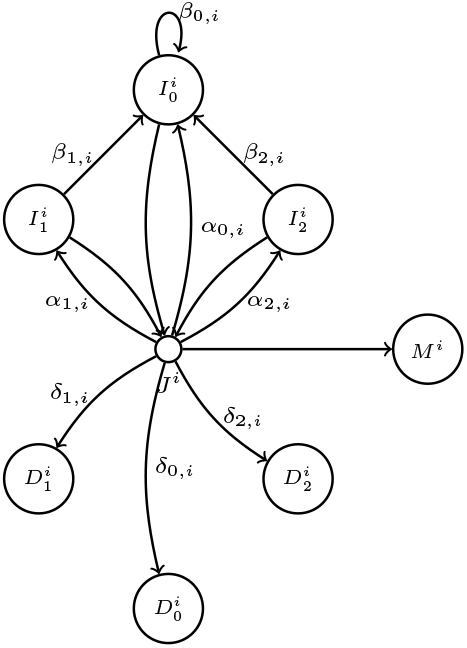
One block of the proposed pHMM structure. See Supplementary Fig. S1 for the full model.

**Figure 2.**
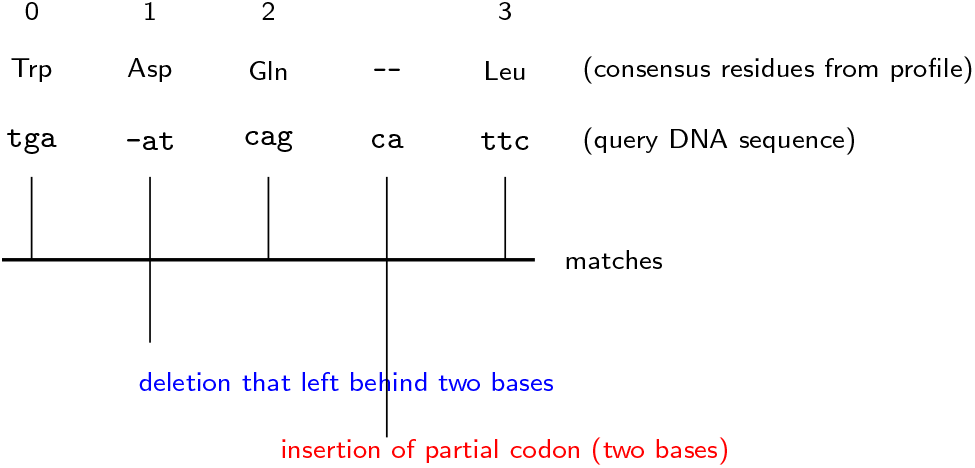
Example alignment.

### Null model

We define a null model that assumes independent and identically distributed codons and allows for frameshift events. Conceptually, it can be viewed as a junction state *Q* with three self-looping emission states: 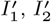, and 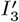, corresponding to single-nucleotide, dinucleotide, and full-codon emissions respectively (see Fig. S2).

A random nucleotide sequence from the null model can be generated by first choosing a reading frame, uniformly at random, each with probability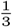. Conditioned on the selected frame, the model generates the sequence sequentially: at each step, the model either generates one of a codon or single-nucleotide or dinucleotide emission, or transitions into a terminal phase. Codons 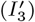 are generated by sampling from an i.i.d. codon distribution, producing three nucleotides at once and preserving the reading-frame structure. Single-nucleotide 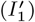 and dinucleotide 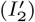 emissions represent frameshift-like events that disrupt the reading frame, produced by drawing from an i.i.d. nucleotide distribution.

The sequence may start and end with partial codons, composed of 1 or 2 i.i.d. single nucleotides.

Given a query sequence, we compute the probability *P* (query seq | null) using the algorithm in Supplementary Material Section S2.

### Scoring

A local alignment *π* is a walk through the profile from left to right that, as it reads the query DNA from left to right, visits a sequence of match states, possibly interleaved with insertions and deletions, consuming the query characters at the different non-silent states. It may be flanked by unaligned ends of the profile and sequence. For a junction state *J*^*i*^ and a position *j* in the query sequence, let

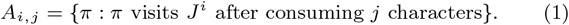

We compute the forward-backward log-likelihood score defined as:

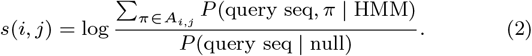

Scores *s*(*i, j*) for all *i, j* are computed by the Forward-Backward Algorithm shown in Supplementary Material Section S3.

High scores indicate strong local correspondences between regions of the query sequence and the profile, and we will use them as seeds to be extended into alignments.

### Significance of score

We compute the number of *i, j* pairs with score *s*(*i, j*) ≥ *s* when comparing a random sequence to our frameshift-ware profile as:

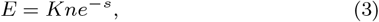

where *n* is the length of the query sequence, and *K* is a Gumbel parameter which can be estimated empirically. Here a random sequence is one generated by the null model, i.e. drawn from the profile’s background codon distribution with frameshift events injected at a configurable probability. Our E-value computation follows from DUMMER which in turn is based on a conjecture by Yu and Hwa [17; 8].

Following DUMMER, the Gumbel parameter *K* is estimated by Monte Carlo simulation. For a profile, we generate random sequences and compute the forward-backward scores. We then fit the scores to a Gumbel distribution via the method of moments to estimate *K*. The estimated parameters are cached to disk avoiding recomputation across runs.

We provide empirical support for the reliability of our E-values.

### Representative alignment construction

Since the score described in Eq. 2 is for a set of alignments, we construct a single representative alignment to be returned to the user. For this, we first rank every sequence position *j* by its best *s*(*i, j*) over *i*, keeping those passing the E-value threshold, and sorting them in decreasing order as alignment seeds. Each seed is then grown outward in both directions using a dynamic programming algorithm that maximizes a score containing contributions from forward-backward likelihood scores of match states and from unaligned null model states. Details are in Supplementary Material Section S4. The space the growing alignment occupies in the sequence is blocked off so that weaker-scoring seeds that try to grow into them are discarded as duplicate alignments. The resulting set of match sequences are presented as gapped alignments to the user.

HMMER3 and BATH have a dedicated multihit feature in which different parts of the query can align to the same profile repeatedly, which is useful in finding repeated domains. While we can construct multiple alignments for a query sequence, our HMM assumes an alignment is only surrounded by unaligned sequences, so there is no favorable scoring for alignments that occur multiple times.

### Speed-up heuristics

We filter reads, or portions thereof, that are unlikely to produce high-scoring alignments against a given profile, in multiple stages.

1. **Ungapped alignment filter** — MMseqs2 first performs a GPU-accelerated ungapped prefilter on the six-frame translation of the query with a configurable p-value threshold defaulting to 0.02.
2. **Gapped, non-frameshift-aware Smith–Waterman filter** — MMseqs2 then runs a gapped Smith–Waterman alignment on the CPU using SIMD acceleration using a configurable, loose P-value threshold (`--prefilter-pvalue`, default 0.005, or 0.02 for increased sensitivity). Surviving hits are merged into contiguous genomic windows and extracted as substrings from the query sequence.
3. **Gapped, frameshift-aware backward-only filter** — We run a SIMD-accelerated backward pass on each extracted substring as detailed in Supplementary Material Section S3.3. E-value of backward scores are computed as in Eq. 3, and substrings with an E-value above the user-configured threshold are discarded.

The full forward-backward algorithm, E-value estimation, and alignment construction, are reserved for the most promising substrings.

### Implementation

Our pHMM is implemented in the software tool DUMMEST (DUMMER without Explicit Sequence Translation). DUMMEST is implemented in C++ and uses Kokkos for vendor-neutral SIMD parallelism. MMseqs2 prefiltering is orchestrated by a Python wrapper that uses file I/O to transfer data between MMseqs2 and DUMMEST without consuming extra memory. A one-time overhead is required for fitting the Gumbel distribution parameters for each protein profile, but subsequent searches are fast. The source code is available at https://github.com/bioinfodlsu/dummest. DUMMEST provides an inter-sequence SIMD implementation that can align 4, 8, or 16 sequences at a time, depending on configured floating-point precision and hardware capabilities. It utilizes Kokkos’ hardware-agnostic Kokkos::SIMD to allow more portability than HMMER3 and BATH’s SIMD implementation. The implementation only makes use of SIMD floating-point arithmetic and shuffling operations, SIMD primitives that are common to most SIMD architectures and thus exposed by Kokkos::SIMD.

Unlike HMMER3 and BATH, which correct for compositional bias by adjusting model parameters on a per-sequence basis, we address it through two mechanisms: masking tandem repeats with TANTAN [6] and setting the background emission probability for each amino acid to be the geometric mean of its position-specific probabilities across the profile following [2] to avoid matches solely due to similar compositions.

Profiles are built with HMMER3’s `hmmbuild` from protein alignments; since `hmmbuild` supports protein only, frameshift background rates default to values estimated from `Badread nanopore2020` model [14] from observed k-mer length changes, and are user-overridable.

## Results

### E-value Reliability

Fig. 3 shows the distributions of the forward-backward scores *s*(*i, j*) and the backward scores used by the filter, for two randomly chosen Pfam profiles of 1000 randomly generated 5000 bp sequences emitted using the profiles’ background codon emission probabilities. They show that the scores for random sequences tend to follow the fitted Gumbel distribution parameters, with a scale of *λ* ≈ 1 as conjectured by [17] and [8] for the non-frameshift case.

**Figure 3.**
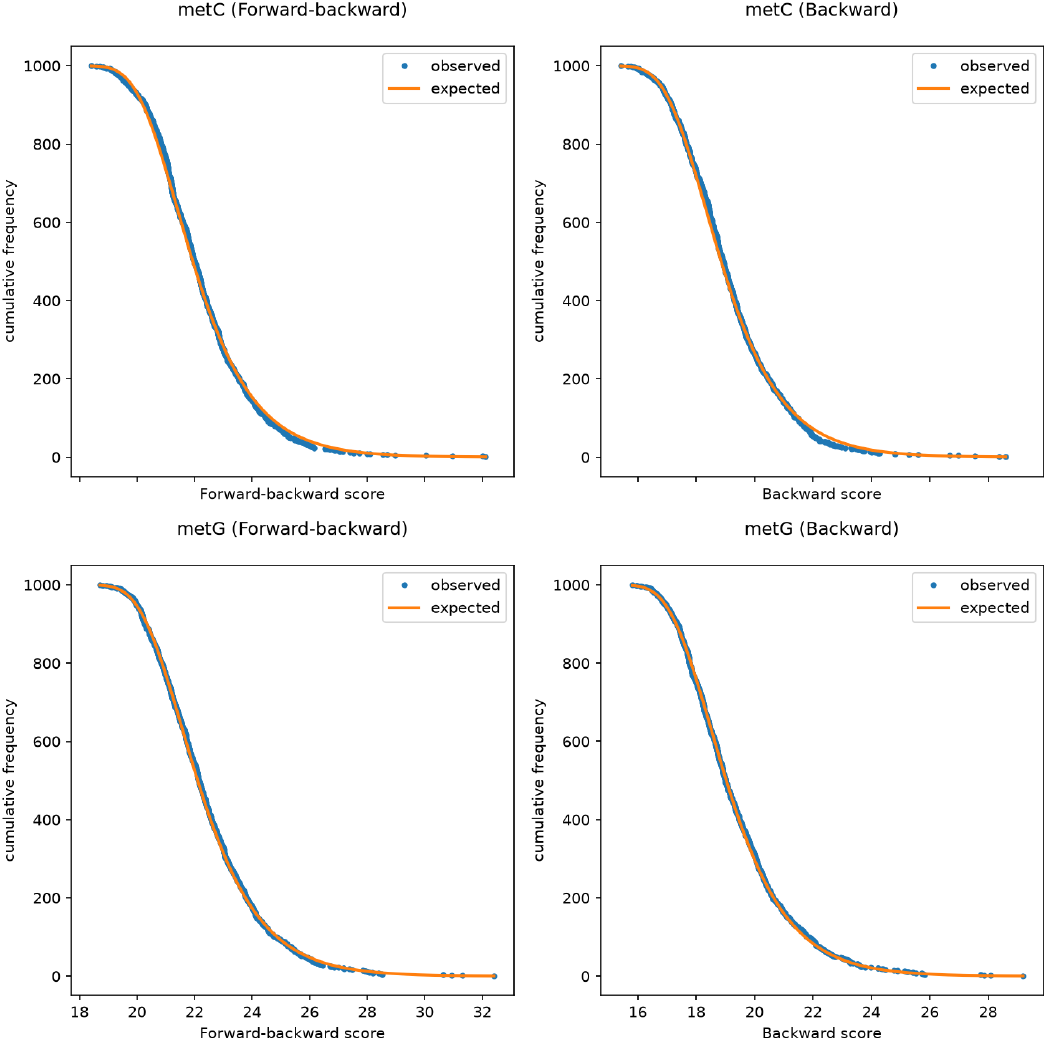
Distribution and fit of random forward-backward score *s*(*i, j*) and backward score used in the filtering.

### Performance on simulated data

We constructed a dataset using transmark [11], modifying it to reflect realistic long-read metagenomics use cases. We randomly selected 1500 Pfam protein families, and divided the sequences of each family into train and test sets, ensuring that no sequence in the test set is more than 50% similar to a sequence in the train set. The 10 “pseudochromosomes,” each approximately 100 Mbp in length, were built from a background sequence emitted from a background DNA HMM, into which the test sequences were embedded. We also planted decoy sequences: randomly chosen protein sequences from the Pfam seed alignments that were shuffled and reverse translated to DNA covering about 5% of each pseudochromosome. The HMM-generated background and the shuffled sequences together serve as the non-homologous regions. Henceforth, embedded sequences comprise both the embedded test sequences and the planted decoy sequences; background denotes all other regions. Simulated nanopore long reads were then generated from these pseudochromosomes using Badread [14] with the `nanopore2020` error model with 5× read depth. Badread also replaced 1% of all reads with low-complexity junk reads and an additional 1% with completely random sequences. In total, 356,582 simulated long reads were generated.

DUMMEST built profiles with `hmmbuild` and BATH (v2.0 release candidate rc5 [1]) with `bathbuild` from the train set. We aligned the simulated long reads to the pHMMs. To evaluate the impact of the MMseqs2-based prefilter on sensitivity, we also tested a more sensitive configuration of DUMMEST with a gapped prefilter p-value cutoff of

0.02 instead of 0.005.

Hits are classified by their overlap with the embedded test and decoy sequences. A hit is a true positive if it overlaps an embedded test sequence of the same family, on the same strand, by more than half the length of the shorter of the two sequences. Among false positives we distinguish decoy hits, which overlap a planted shuffled decoy sequence; background hits, which overlap neither a test sequence nor a decoy sequence and arise from the background sequence, junk and random synthetic reads; and wrong-target hits, which overlap an embedded test sequence but are assigned to a different family and/or the opposite strand. Decoy, true-positive and wrong-target hits are deduplicated per embedded sequence within each hit category, keeping the smallest E-value per sequence per category, to avoid inflating counts from multiple fragmented alignments to the same locus. Background hits have no clear identity for deduplication and are therefore reported as raw counts.

Fig. 4 shows for varying E-value thresholds, the proportion of embedded sequences recovered and the number of false positive hits divided by the number of profiles. DUMMEST found more true positive hits at low false positive thresholds than BATH. At a relatively stringent E-value threshold of 10^−5^, BATH found more hits in general (42,280 vs. 39,447) but also produced more decoy false positives (Table 1). DUMMEST’s single background hit (E-value 4.3 × 10^−6^ in Chloroa b-bind) overlaps a valid alignment but overextends into non-homologous sequence so that its overlap falls below the 50% threshold required for a true positive. For BATH, its most significant decoy hit had an E-value of 9.4×10^−12^, a poly-asparagine repeat in Secretin N 2 that aligned despite the use of a bias filter. DUMMEST avoided that scenario by masking tandem repeats in the profile. Both tools also produced about 120 wrong-target false positives at 10^−5^ (Table 1), comprising hits assigned to a different family than the embedded target or reported on the opposite strand. These may partly reflect distant homology or domains shared between Pfam families.

**Table 1.** Alignment performance on simulated metagenomic reads at E-value threshold of 10^−5^. Mean coverage is the mean percentage of embedded nucleotides recovered per true-positive hit; mean overextension is the mean number of nucleotides extending into non-homologous regions. False positives are split into decoy hits, background hits and wrong-target hits. Decoy, true-positive and wrong-target counts are deduplicated per embedded sequence (lowest E-value kept); background hits are reported as raw counts.

| Tool | True-positive hits | False-positive hits |  |  | Mean coverage (%) | Mean overextension (nt) |
| --- | --- | --- | --- | --- | --- | --- |
|  |  | decoy hits (non-target) | background hits (non-target) | wrong-target |  |  |
| BATH | 42280 | 90 | 0 | 122 | 89.14 | 0.16 |
| BATH (non-frameshift-aware) | 11488 | 0 | 0 | 35 | 46.15 | 0.03 |
| DUMMEST | 39447 | 0 | 1 | 120 | 90.29 | 0.09 |
| DUMMEST (more sensitive) | 40948 | 0 | 1 | 129 | 90.11 | 0.09 |

**Figure 4.**
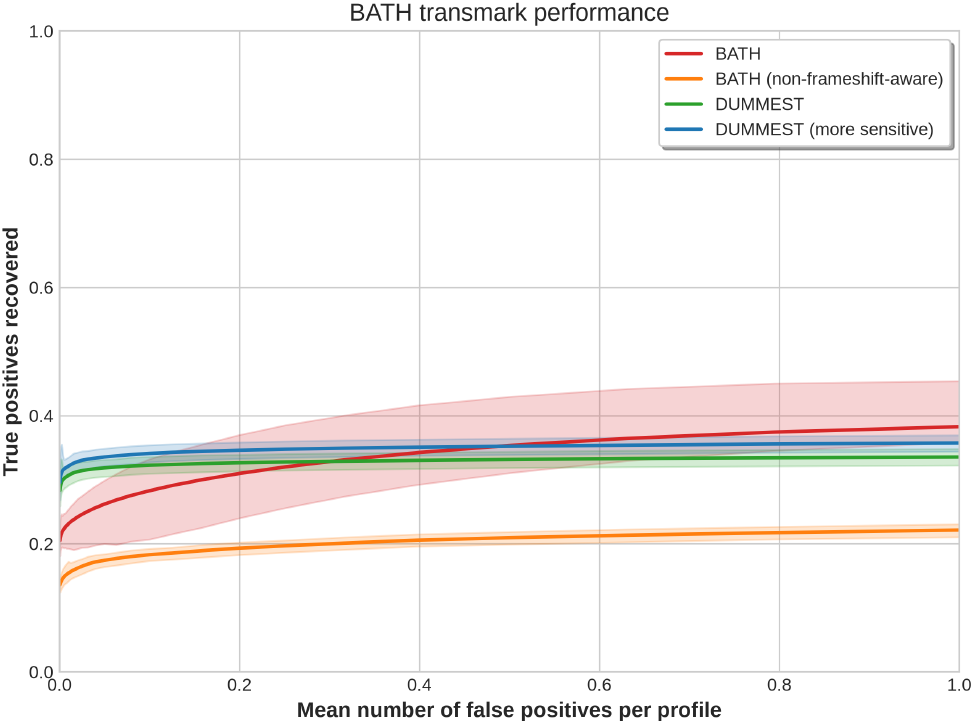
True positives vs. mean number of false positives per profile for simulated metagenomic reads.

Apart from evaluating performance based on the number of hits, we assessed alignment coverage as the percentage of the embedded nucleotides recovered by the true positive hits and alignment overextension as the mean number of nucleotides extending into non-homologous regions. Table 1 shows that DUMMEST and BATH have similar coverage (90.29% and 89.14%) and overextension (0.09 and 0.16 nucleotides). It indicates that DUMMEST’s alignments are not less precise at their boundaries despite its simpler model.

Among true and false positives, Fig. 5 shows that DUMMEST and BATH E-values were heavily correlated, providing additional evidence that DUMMEST’s E-value estimates are sensible. Most high-scoring hits receive higher E-values from DUMMEST, the opposite of what was observed for DUMMER when compared against nhmmer [7]. Because they must handle frameshifts, DUMMEST and BATH both use modified null models relative to their progenitors DUMMER and HMMER3, respectively. Unlike protein-to-protein and nucleotide-to-nucleotide alignment, no studies have compared null models for frameshift-aware alignment. Future work could examine the effects of different null models (e.g. nucleotide-only, reverse-sequence) in the frameshift-aware case.

**Figure 5.**
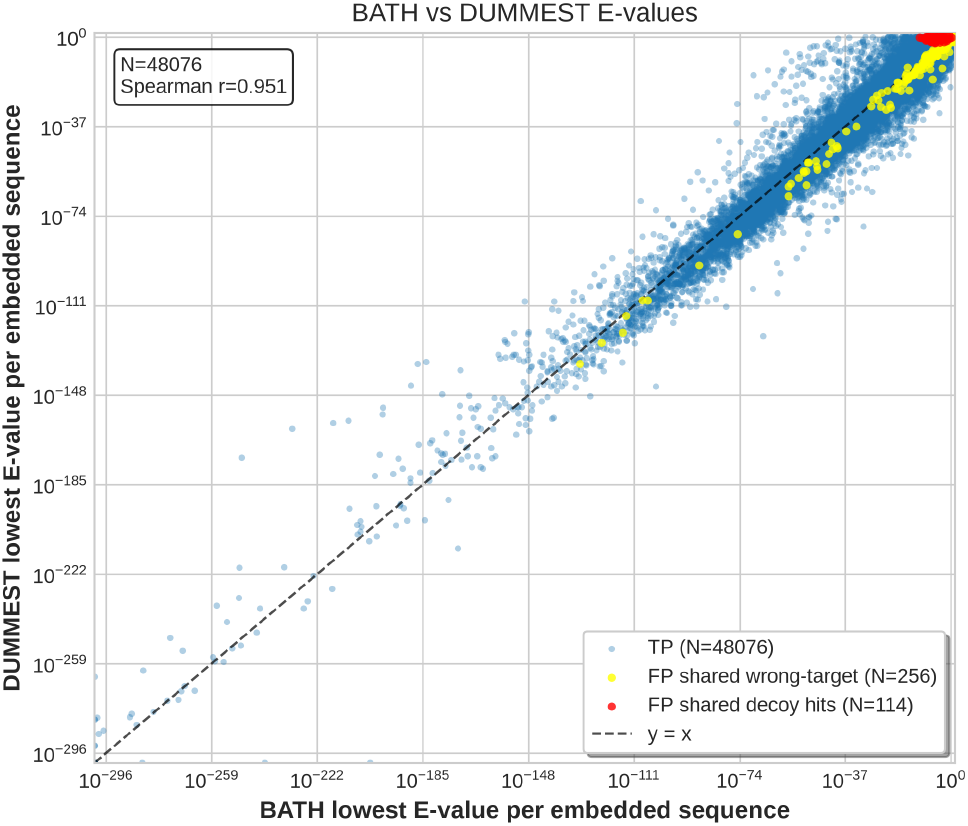
Comparison of DUMMEST and BATH E-values for true and false positives. Lowest E-value per embedded sequence and hit category (one point per sequence per category).

Among hits present only in BATH, a majority of them were in the twilight zone (close to 10^−5^). Unlike BATH’s Viterbi pre-filter, the MMseqs2 gapped pre-filter does not feature position-specific insertion and deletion probabilities, possibly hindering its sensitivity. Furthermore, the frameshift parameters are approximated from the error model. An approach obtaining frameshift parameters from the query sequence itself, like last-train [15], could yield better-fitting parameters to compensate for the simpler model of DUMMEST, which could be less forgiving to errors.

### Performance on Real Data

To verify performance in real-life data, we performed a case study to identify antibiotic resistance genes (ARGs) in Oxford Nanopore long reads collected from wastewater influent (NCBI SRA SRR12664565) [12], using reference protein families from AMRFinderPlus [5]. To be able to compare against non-frameshift-aware methods, the long reads were polished with Ratatosk [9] using paired-end Illumina short reads produced by the same study. In total, there were 2.3 million real long reads, with a total length of approximately 5 Gbp.

In the absence of a ground truth, we measured performance using two proxy metrics: (i) the total number of unique (read id, protein family) pairs, and (ii) the cumulative alignment coverage measured in base pairs.

Fig. 6 and Table 2 show performance in real data at *E* ≤ 1. DUMMEST is generally comparable to BATH, noticeably outperforming it compared to the previous benchmark. At a fixed E-value cutoff of 1 × 10^−5^, BATH and DUMMEST are frequently in agreement, sharing 49,077 pairs with a Jaccard index of 0.67 (3,466 only in BATH and 20,202 only in DUMMEST; union 72,745). Despite the usage of polishing, both tools outperform HMMER3 and BATH frameshift turned off (–no-fs), suggesting that polishing is limited in its ability to remove frameshift-inducing indels.

**Table 2.**
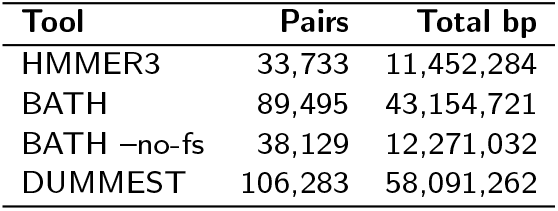
Comparison of tools by number of (read id, profile) pairs and total base pairs at E-value threshold *E* ≤ 1.

**Figure 6.**
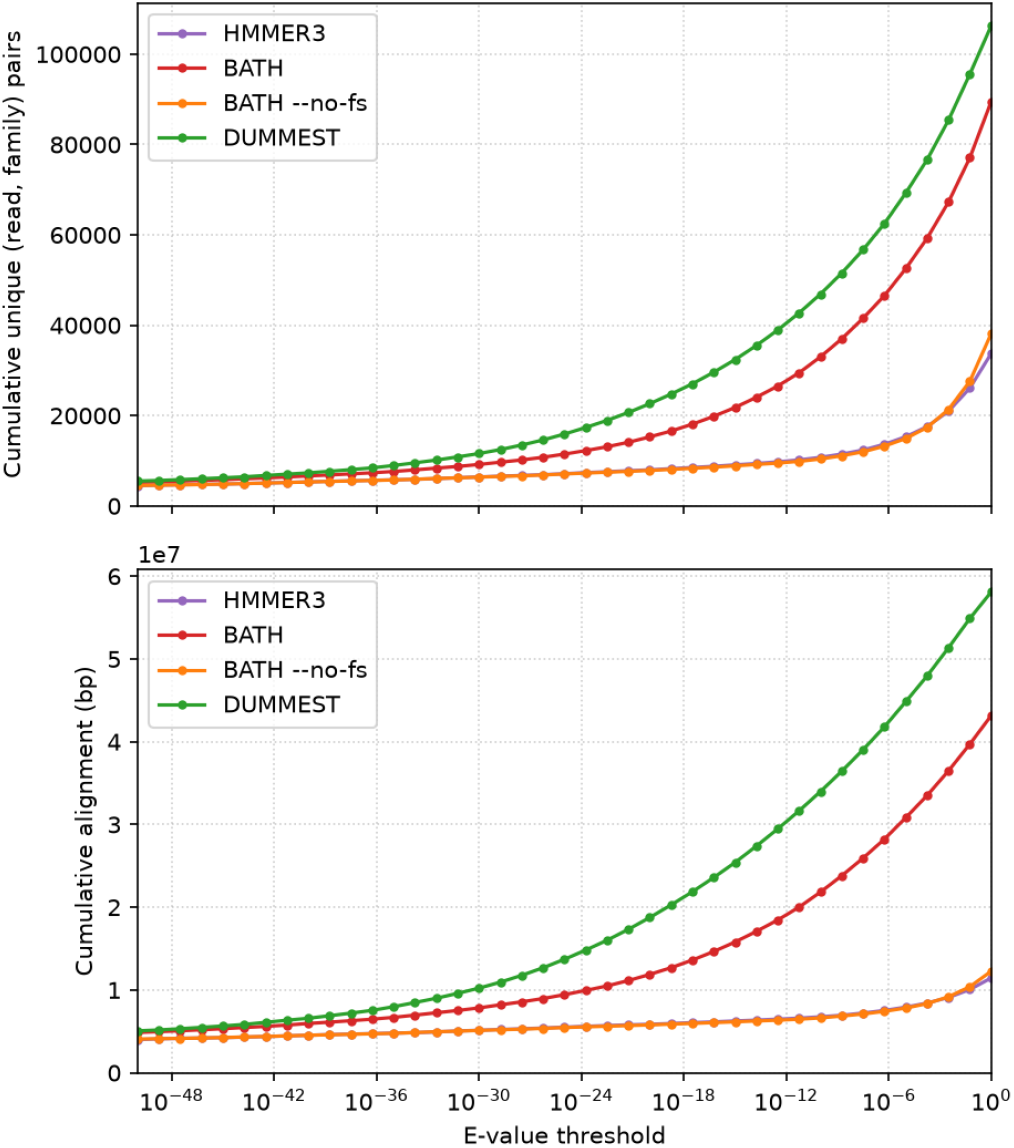
Cumulative number of hits and alignment coverage length by E-value threshold. Table 2 reports values at *E ≤* 1.

To further verify the specificity of DUMMEST, we tested its ability to find ARGs on the complete genome sequence of *Pandoravirus salinus*, as viruses typically lack ARGs. When running with the ungapped prefilter disabled, it found one significant hit for trimethoprim resistance (*E* ≈ 7 × 10^−11^). This is expected, as Pandoraviruses are known to carry trimethoprim resistance genes [16]. No other gene in the profile database used is known to be carried by Pandoraviruses.

### Runtime analysis

Table 3 shows the runtime on both datasets. For the experiments which used the prefilter, we used the full datasets, which have approximately 5 Gbp of query sequences and 1, 500 profiles for the transmark dataset and 5 Gbp and 143 profiles for the wastewater dataset.

**Table 3.** Wall-clock time. 8 threads / 32 GB RAM / RTX 5070 Mobile GPU.

| Tool | Dataset | Wall (s) | Prefilter |
| --- | --- | --- | --- |
| DUMMEST | TRANSMARK | 1891.5 | on |
| DUMMEST (MORE SENSITIVE) | TRANSMARK | 2344.3 | on |
| BATH | TRANSMARK | 22095.0 | on |
| BATH -NOFS | TRANSMARK | 21446.0 | on |
| DUMMEST | wastewater | 1033.8 | on |
| BATH | wastewater | 2433.0 | on |
| BATH -NOFS | wastewater | 2219.3 | on |
| HMMER3 | wastewater (1000-sequence subset) | 3158.1 | on |
| DUMMEST | wastewater (1000-sequence subset) | 385.5 | off |
| BATH | wastewater (1000-sequence subset) | 1136.5 | off |

Six-frame translation and MMseqs2 database construction time are excluded for DUMMEST and HMMER3. Using an SSD, these operations took six minutes for the wastewater dataset, while it took one minute in transmark. Six-frame translation is bottlenecked by seqtk’s use of a single thread to decompress gzip-compressed sequence files during the evaluation, which is avoidable in practice by decompressing with other tools.

The prefilter-less experiments show that DUMMEST’s speedup is not merely from the use of GPU-accelerated prefiltering: the full forward-backward implementation is 3× faster than BATH. DUMMEST’s runtime advantage derives from two factors: (i) a simplified pHMM model, and (ii) GPU-accelerated pre-filtering via MMseqs2, which eliminates non-homologous query-target pairs faster than BATH.

The speedup in the wastewater dataset is noticeably less than in transmark. It could be caused by the high number of ARG hits in wastewater influent, as BATH and DUMMEST identified around 89, 000–106, 000 pairs, compared to around 40, 000 true positives in transmark. No amount of filtering can speed up a dataset that has such a dense concentration of hits. While the initial release of BATH tried to avoid running the frameshift-aware alignment as it was originally slow, such optimization will probably not cause a huge speedup as non-frameshift-aware tools could only identify less than half of BATH and DUMMEST’s hits here. A promising approach is to calculate only the high-scoring cells of the forward-backward matrix, as in nail [13]. In our initial experiments with that, it appears that MMseqs2 may not provide good enough locations of high-scoring regions to make this work practice, although it was not fully investigated. The backward-only filter is another promising source of seeds for future iterations, as it is more sensitive than any Viterbi or Smith-Waterman-based filter. There is also further headroom from engineering improvements in the current implementation, as the null-model precomputation is not yet fully SIMD-accelerated.

## Conclusion

We have presented DUMMEST, a simpler frameshift-aware pHMM based on the DUMMER architecture. By extending DUMMER’s match, insertion, and deletion states to emit nucleotides and partial codons, DUMMEST models frameshift events in a simpler manner. Empirical benchmarks show that DUMMEST is comparable to BATH in sensitivity and alignment coverage, shows better specificity when sequencing artifacts are introduced into simulated genomes with known ground truth, while running 2–12× faster depending on dataset (∼11.7× on transmark and ∼2.4× on wastewater; ∼2.9× for the prefilter-less core alignment), and produces alignment scores that likely follow the universal E-value statistics of Yu and Hwa [17].

Several modeling directions remain open. No studies have compared null models for frameshift-aware alignment, so alternatives such as nucleotide-only or reverse-sequence null models could be examined. Frameshift parameters are currently approximated from the error model; estimating them from the query sequence itself, as last-train does, could further improve sensitivity.

On the engineering side, the main limitations are DUMMEST’s high memory requirements, due to its need to store sequences in memory, and its I/O usage, driven by the use of large temporary files to pass filtered sequences from MMseqs2 to DUMMEST. It also lacks a customized method for building HMMs and instead relies on HMMER’s `hmmbuild`. Future work could address these limitations by streaming sequences in batches and speeding up score estimation, for example, by computing only the high-scoring cells of the forward-backward matrix, as in nail [13].

## Acknowledgements

We thank Martin Frith for insightful discussions.

H.S.C. would like to express his gratitude to the Philippine Department of Science and Technology – Science Education Institute (DOST-SEI) for the financial support provided through the Engineering Research and Development for Technology (ERDT) Graduate Scholarship Program.

