## Supplementary Material for "DUMMEST: A simpler and faster model for frameshift-aware alignment of a DNA sequence to a protein family"

#### S1 Model specification

##### S1.1 States

The full model is shown in Fig. S1. The gray states represent the background or unaligned flanks of the query, where there is no homology to the profile, while the dark states represent the protein family.

An alignment begins at the Begin state. A query DNA sequence has three possible reading-frame offsets in the given sequence orientation, chosen with probabilities $\rho_0$, $\rho_1$, $\rho_2$ (1/3 each at default). $V_0$ emits the initial full codon, $V_1$ and $V_2$ emit the last one and two nucleotides of the initial codons, respectively, so that subsequent codons are in frame. Gray state $Q_b$ is the null model junction at the beginning of the profile, which allows emitting background or unaligned sequence via three insertion states: single-nucleotide ($I'_1$), dinucleotide ($I'_2$), and codon ($I'_3$). From the unaligned upper flank—the chain of $U^i$ states—the model dips into any block through $U^i \to J^i$, where $i = 0, \ldots, m$ indexes the junctions between profile positions, $m$ is the profile length (including the extra junction $J^m$ after the last match for post-profile insertions), and the $J^i$ are the silent junctions connecting them.

Each block $i$ in the aligned region contains a match state $M^i$, three insertion states $I^i_0$, $I^i_1$, $I^i_2$, and three deletion states $D^i_0$, $D^i_1$, $D^i_2$. The match state $M^i$ emits a codon whose probability is determined by the amino acid at that profile position. The insertion states model additional nucleotides relative to the profile: $I^i_0$ emits a full codon, while $I^i_1$ and $I^i_2$ emit one and two nucleotides, respectively, corresponding to frameshifting insertion events. The deletion states allow skipping profile positions: $D^i_0$ skips a full codon, while $D^i_1$ and $D^i_2$ emit two and one nucleotides respectively, accounting for DNA deletions that are not multiples of three.

Exiting, each block connects back to the unaligned lower flank—the chain of $L^i$ states—via $J^i \to L^i$, and $L^m$ leads to the null model junction $Q_e$ at the end of the profile, which reuses the same insertion states ($I'_1$, $I'_2$, $I'_3$) for emitting in the background or unaligned sequence before reaching the End state.

To summarize, the model is comprised of the following states:

| | |
|---|---|
| Begin | : silent start state |
| $V_0, V_1, V_2$ | : initial-codon frame-selection states |
| $Q_b$ | : silent junction at the entry flank of the profile |
| $I'_1, I'_2, I'_3$ | : insertion states in the unaligned flank (same states reused at both flanks) |
| $U^i$ | : silent flank junction above block $i$ (small gray circle in Figure S1) |
| $J^i$ | : silent junction states connecting profile positions (small black circles in Figure S1) |
| $M^i$ | : codon match |
| $I^i_0, I^i_1, I^i_2$ | : insert |
| $D^i_0, D^i_1, D^i_2$ | : delete |
| $L^i$ | : silent flank junction below block $i$ (small gray circle in Figure S1) |
| $Q_e$ | : silent junction at the exit flank of the profile |
| End | : silent end state |

##### S1.2 Emission probabilities

Emissions use the following distributions. Codons are written as $b_1b_2b_3$ in emission order.

$\psi(y)$ : codon distribution over length-3 strings $y = b_1b_2b_3$ (emission order)
$\phi(b)$ : per-base distribution over single DNA bases, used for partial-codon emissions
$\theta_i(y)$ : match-state codon emission at profile position $i$

For partial-codon emissions the emitted nucleotides are the first one ($b_1$) or first two ($b_1b_2$) in emission order. A state emitting three nucleotides at once uses the codon distribution ($\theta_i$ for match states, $\psi$ for insertion states); a state emitting one or two nucleotides multiplies the per-base $\phi$ once per emitted base.

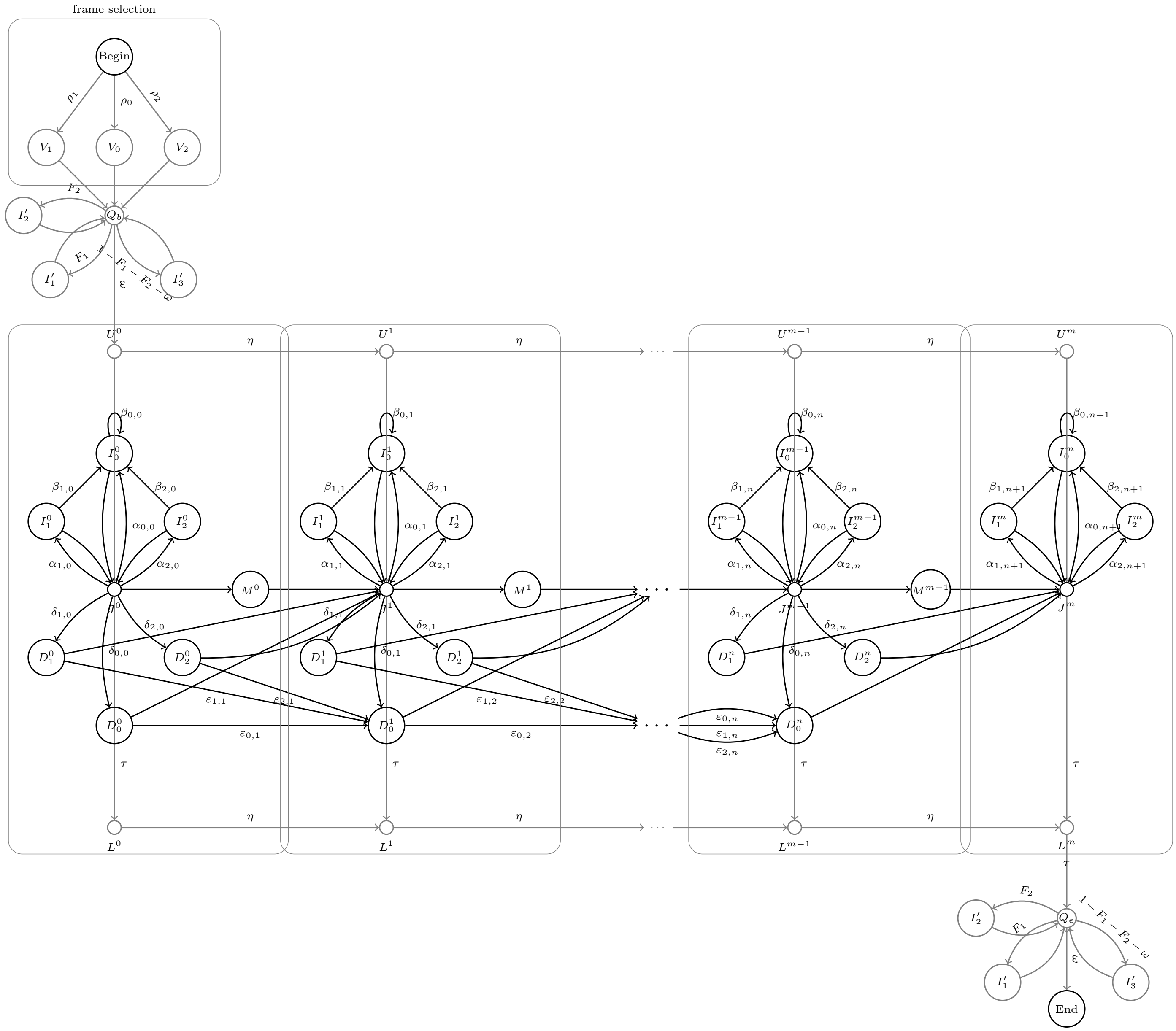


Figure S1: Profile HMM structure.

| State | Emission length (nt) | Emission prob. of the emitted string |
|---|---|---|
| $V_0$ | 3 | $\psi(b_1b_2b_3)$ |
| $V_1$ | 1 | $\phi(b_1)$ |
| $V_2$ | 2 | $\phi(b_1)\phi(b_2)$ |
| $I'_1$ | 1 | $\phi(b_1)$ |
| $I'_2$ | 2 | $\phi(b_1)\phi(b_2)$ |
| $I'_3$ | 3 | $\psi(b_1b_2b_3)$ |
| $M^i$ | 3 | $\theta_i(b_1b_2b_3)$ |
| $I^i_0$ | 3 | $\psi(b_1b_2b_3)$ |
| $I^i_1$ | 1 | $\phi(b_1)$ |
| $I^i_2$ | 2 | $\phi(b_1)\phi(b_2)$ |
| $D^i_0$ | 0 | – |
| $D^i_1$ | 2 | $\phi(b_1)\phi(b_2)$ |
| $D^i_2$ | 1 | $\phi(b_1)$ |

#### S1.3 Transition probabilities

We use the following notations for the transition probabilities.

$\rho_f$ : Begin $\to V_f$, $f \in \{0,1,2\}$ (choose reading frame)
$\eta : U^i \to U^{i+1}$, $L^i \to L^{i+1}$ (remain in the unaligned flank)
$\omega : Q_b \to U^0$ (enter an alignment); $Q_e \to$ End (end the HMM)
$F_1 : Q \to I'_1$, $Q \in \{Q_b, Q_e\}$ (single-nucleotide emission in the unaligned flank)
$F_2 : Q \to I'_2$, $Q \in \{Q_b, Q_e\}$ (dinucleotide emission in the unaligned flank)
$\tau : J^i \to L^i$; $L^m \to Q_e$ (exit an alignment)
$\alpha_{0,i} : J^i \to I^i_0$ (start a full-codon insertion)
$\alpha_{1,i} : J^i \to I^i_1$ (start a single-nucleotide insertion)
$\alpha_{2,i} : J^i \to I^i_2$ (start a dinucleotide insertion)
$\beta_{0,i} : I^i_0 \to I^i_0$ (extend a full-codon insertion)
$\beta_{1,i} : I^i_1 \to I^i_0$ (return from a single-nucleotide insertion to a full codon)
$\beta_{2,i} : I^i_2 \to I^i_0$ (return from a dinucleotide insertion to a full codon)
$\delta_{0,i} : J^i \to D^i_0$ (start a deletion)
$\delta_{1,i} : J^i \to D^i_1$ (start a frameshifted deletion)
$\delta_{2,i} : J^i \to D^i_2$ (start a frameshifted deletion)
$\varepsilon_{0,i} : D^{i-1}_0 \to D^i_0$ (extend a deletion)
$\varepsilon_{1,i} : D^{i-1}_1 \to D^i_0$ (extend a frameshifted deletion)
$\varepsilon_{2,i} : D^{i-1}_2 \to D^i_0$ (extend a frameshifted deletion)

The remaining profile transitions that are not the only out-edge are obtained as complements of the probabilities above.

$1 - F_1 - F_2 - \omega : Q \to I'_3$, $Q \in \{Q_b, Q_e\}$ (codon emission in the unaligned flank)
$1 - \eta : U^i \to J^i$ (enter an alignment block from the unaligned flank)
$1 - \sum_k \alpha_{k,i} - \sum_k \delta_{k,i} - \tau : J^i \to M^i$ (enter a match state)
$1 - \beta_{k,i} : I^i_k \to J^i$, $k \in \{0,1,2\}$ (end an insertion sequence)
$1 - \varepsilon_{k,i+1} : D^i_k \to J^{i+1}$, $k \in \{0,1,2\}$ (end a deletion sequence)

The probability of remaining in the unaligned flank ($\eta$) is near 1, while the probabilities of transitioning into an alignment ($\omega$), out of an alignment ($\tau$), and to the end state ($\omega$) are near 0, giving the model no alignment-length bias [1]. The ~~symbols~~ parameters $\omega$ and $\tau$ are each shared between entry and exit transitions. From $Q_e$, the model transitions to the silent End state with probability $\omega$, terminating the alignment.

#### S1.4 Setting/training of parameters

##### S1.4.1 Conversion of HMMER3 insertion/deletion probabilities

Protein profiles can be generated by either HMMER's `hmmbuild` or DUMMER's `hmmbuild`, although HMMER's `hmmbuild` is recommended because it is more mature. The current implementation obtains $\alpha_{0,i}$, $\beta_{0,i}$, $\delta_{0,i}$, $\varepsilon_{0,i}$, and $\psi(y)$ from HMMER profile files using the same method described in [1].

For a length-$m$ profile built by HMMER3, the profile file provides the following quantities for $0 \le i \le m$:

$M^i \to I^i$: probability of starting an insertion at position $i$,

$I^i \to I^i$: probability of extending an insertion at position $i$,

$M^i \to D^{i+1}$: probability of starting a deletion after position $i$,

$D^i \to D^{i+1}$: probability of extending a deletion from position $i$.

The transition probabilities $D^0 \to D^1$, $M^m \to D^{m+1}$, and $D^m \to D^{m+1}$ are always zero. HMMER profile files are 1-indexed, with $e_i(y)$, the emission probability of amino acid $y$ at position $i$, for $0 < i \leq m$ (HMMER's own 1-indexed notation, hence subscript); the mapping below converts them to our 0-indexed parameters. HMMER3 does not account for frameshifts in its transitions, so we add $\alpha_{1,i}$, $\alpha_{2,i}$, $\delta_{1,i}$, $\delta_{2,i}$ and rescale the HMMER insertion and deletion starts by $(1 - \alpha_{1,i} - \alpha_{2,i} - \delta_{1,i} - \delta_{2,i})$ to leave room for frameshift starts:

These are mapped to our non-frameshifted transition parameters as follows:

$$\alpha_{0,i} = (M^i \to I^i)(1 - \alpha_{1,i} - \alpha_{2,i} - \delta_{1,i} - \delta_{2,i}) \qquad \text{for } 0 \leq i \leq m \tag{1}$$

$$\beta_{0,i} = (I^i \to I^i) \qquad \text{for } 0 \leq i \leq m \tag{2}$$

$$\delta_{0,i} = (M^i \to D^{i+1})(1 - \alpha_{1,i} - \alpha_{2,i} - \delta_{1,i} - \delta_{2,i}) \qquad \text{for } 0 \leq i < m \tag{3}$$

$$\varepsilon_{0,i} = (D^i \to D^{i+1}) \qquad \text{for } 0 < i < m \tag{4}$$

##### S1.4.2 Frameshifted insertions/deletions probabilities

Frameshifted insertion ($\alpha_{1,i}$, $\alpha_{2,i}$) and deletion ($\delta_{1,i}$, $\delta_{2,i}$) probabilities are position-independent in the current implementation. The specific values used are extracted from the `Badread` `nanopore2020` error model. The unaligned flank's frameshift transition probabilities are then set as: $F_1 = \alpha_{1,i} + \delta_{2,i}$ and $F_2 = \alpha_{2,i} + \delta_{1,i}$, which combine insertion and deletion events that have the same net effect on the reading frame.

Currently, the extraction script `tools/get_indel_rates.py` estimates indel proportions from the table of k-mer alternatives for each possible 7-mer in the `Badread` model. Its output is an occurrence rate per reference base, not a probability of an HMM transition. This will be replaced with either a simulator-generic approach that generates simulated reads, and tries to fit the indel proportions of the HMM using expectation-maximization or a `last-train`-like approach that estimates frameshift rates from the query sequences themselves.

The start transition probabilities for frameshifted insertions and deletions are set directly to these values obtained from the script, independent of profile position:

$$\alpha_{1,i} = 0.0171, \qquad \alpha_{2,i} = 0.0018. \tag{5}$$

$$\delta_{1,i} = 0.0328, \qquad \delta_{2,i} = 0.0083. \tag{6}$$

In the current implementation, $\beta_{1,i}$, $\beta_{2,i}$, $\varepsilon_{1,i}$, and $\varepsilon_{2,i}$ are set to zero, meaning frameshifted insertions and deletions do not chain. The parameters are retained in the model description for potential future use.

##### S1.4.3 Emission probabilities from protein profiles

A difference from HMMER and DUMMER is that stop codons are handled probabilistically, as stop codons could be affected by frameshifts. In the current implementation, the overall probability of emitting a stop codon is hardcoded as $S := 0.0005$.

The emission probability $\theta_i(y)$ for a sense codon $y$ is derived from the profile HMM's amino acid emission probabilities $e_{i+1}$. Let $aa(y)$ denote the amino acid translated from codon $y$, and let $N_{aa(y)}$ be the number of synonymous codons encoding $aa(y)$ under the standard genetic code. To account for codon degeneracy and reserved stop-codon probability, the codon emission probability is given by:

$$\theta_i(y) = \frac{e_{i+1}(aa(y)) \cdot (1 - S)}{N_{aa(y)}}. \tag{7}$$

For stop codons, $\theta_i$(a stop codon) is defined as $S$ / #stop codons, i.e. 0.0005/3 in our case.

Future work may improve this model by incorporating codon usage information to handle codon degeneracy more precisely than the current uniform distribution.

Let $S_{\text{bg}} := \psi(\text{TAA}) + \psi(\text{TAG}) + \psi(\text{TGA})$ be the total probability of the null model emitting a stop codon. The probability of the null model emitting a sense codon $\psi(y)$, used in insertion states, is the geometric mean of $\theta_i(y)$ over the $m$ match positions ($0 \leq i < m$), as used in [1], scaled by $(1 - S_{\text{bg}})$ to reserve stop-codon probabilities ($S_{\text{bg}} = 3/64$ at default). For stop codons, $\psi$(a stop codon) $= S_{\text{bg}}/3$, i.e. 1/64. We refer to these null-model emission probabilities as background probabilities.

$\phi(b)$, used in the insertion of individual DNA bases, is set uniformly to 0.25 for all bases. The frame-choice probabilities $\rho_f$ are fixed at 1/3 for all $f$, so each frame offset has a uniform probability.

### S2 Null model

The null model is shown in Fig. S2. It contains a silent junction state $Q$ with $V_0/V_1/V_2$ entry states and $I'_1/I'_2/I'_3$ self-loops.

Given a query sequence $s = s_0 s_1 \dots s_{n-1}$ of $n$ nucleotides, where $s_{a..b}$ denotes the inclusive substring from position $a$ to $b$ and query positions satisfy $0 \le j < n$, we wish to compute $P(s \mid \text{null})$. Let $P(s_{0..j} \mid \text{null})$ denote the probability that the null model emits the prefix $s_{0..j}$ (the sum over all null-model paths in this extracted null-only tiling through $V$ and $I'$ states, as opposed to full-model paths that also visit $M/D$ states), and $P(s_{j..n-1} \mid \text{null})$ the analogous probability for the suffix $s_{j..n-1}$.

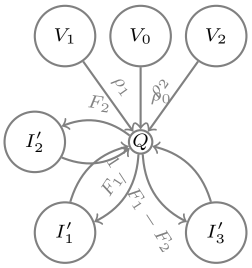


Figure S2: Null model. Prefix paths emit the initial codon with $V_0$ (3 nt, $\rho_0$), $V_1$ (1 nt, $\rho_1$) or $V_2$ (2 nt, $\rho_2$) and tile the remainder with $I'_1$ (1 nt, $F_1$), $I'_2$ (2 nt, $F_2$) and $I'_3$ (codon, $1 - F_1 - F_2$) emissions; $P(s_{0..j} \mid \text{null})$ sums all possible tilings. Truncated suffix emissions at the end are emitted with the probability of each individual nucleotide.

The probability $P(s_{0..j} \mid \text{null})$ is:

$$\begin{aligned} P(s_{0..j} \mid \text{null}) = {} & (1 - F_1 - F_2)\, \psi(s_{j-2..j})\, P(s_{0..j-3} \mid \text{null}) \\ & + F_1\, \phi(s_j)\, P(s_{0..j-1} \mid \text{null}) \\ & + F_2\, \phi(s_{j-1})\, \phi(s_j)\, P(s_{0..j-2} \mid \text{null}), \qquad j \ge 3, \end{aligned} \tag{8}$$

with base cases

$$P(s_{0..j} \mid \text{null}) = \begin{cases} \rho_0\, \psi(s_{0..2}) + F_1\, \phi(s_2)\, P(s_{0..1} \mid \text{null}) + F_2\, \phi(s_1)\, \phi(s_2)\, P(s_{0..0} \mid \text{null}), & j = 2, \\ \rho_2\, \phi(s_0)\, \phi(s_1) + F_1\, \phi(s_1)\, P(s_{0..0} \mid \text{null}), & j = 1, \\ \rho_1\, \phi(s_0), & j = 0. \end{cases} \tag{9}$$

The probability $P(s_{j..n-1} \mid \text{null})$ of generating the suffix $s_{j..n-1}$ is computed via a reverse pass:

$$\begin{aligned} P(s_{j..n-1} \mid \text{null}) = {} & (1 - F_1 - F_2)\, \psi(s_{j..j+2})\, P(s_{j+3..n-1} \mid \text{null}) \\ & + F_1\, \phi(s_j)\, P(s_{j+1..n-1} \mid \text{null}) \\ & + F_2\, \phi(s_j)\, \phi(s_{j+1})\, P(s_{j+2..n-1} \mid \text{null}), \qquad j \le n-3, \end{aligned} \tag{10}$$

with boundary cases

$$P(s_{j..n-1} \mid \text{null}) = \begin{cases} (1 - F_1 - F_2)\, \phi(s_{n-2})\, \phi(s_{n-1}) + F_1\, \phi(s_{n-2})\, P(s_{n-1..n-1} \mid \text{null}) + F_2\, \phi(s_{n-2})\, \phi(s_{n-1}), & j = n-2, \\ (1 - F_1 - F_2)\, \phi(s_{n-1}) + F_1\, \phi(s_{n-1}) + F_2\, \phi(s_{n-1}), & j = n-1, \\ 1, & j = n, \end{cases} \tag{11}$$

They are computed via dynamic programming, using $F_1 = \alpha_{1,i} + \delta_{2,i}$ and $F_2 = \alpha_{2,i} + \delta_{1,i}$ for the background frameshift rates.

The total null probability is then:

$$P(\text{seq} \mid \text{null}) = P(s_{0..n-1} \mid \text{null}) + (1 - F_1)\, \phi(s_{n-1})\, P(s_{0..n-2} \mid \text{null}) + (1 - F_1 - F_2)\, \phi(s_{n-2})\, \phi(s_{n-1})\, P(s_{0..n-3} \mid \text{null}). \tag{12}$$

We employ a probability-ratio trick to directly compute the probability ratio without logarithm operations. The idea is to split the overall null probability into $n$ instances of $\frac{1}{\sqrt[n]{P(s|\text{null})}}$ that is multiplied into the recurrence every time a nucleotide is read. Since each transition that increases the number of nucleotides read performs a multiplication operation, this results in the denominator eventually becoming $P(\text{seq}|\text{null})$, keeping the computation in linear probability space and enabling efficient SIMD arithmetic. This has been observed to be numerically stable for sequences up to approximately 10,000 bases (typical long-read lengths). The implementation carries these values in a custom floating point datatype with a 40-bit exponent field.

The implementation precomputes normalized prefix and suffix arrays $\text{null\_model\_prefix}[j] = P(s_{0..j} \mid \text{null}) \cdot P(s \mid \text{null})^{-(j+1)/n}$ and $\text{null\_model\_suffix}[j] = P(s_{j..n-1} \mid \text{null}) \cdot P(s \mid \text{null})^{-(n-j)/n}$, so that the forward and backward recurrences below need only add the corresponding array entries without additional multiplications.

### S3 Forward–Backward algorithm

#### S3.1 Notation

Let $S_j$ denote the event that a path visits state $S$ after consuming $j$ characters (i.e. after emitting the prefix $s_{0..j}$). In particular, $J_j^i$ and $M_j^i$ denote visits to junction $J^i$ and to match $M^i$ while emitting the codon $s_{j-2}s_{j-1}s_j$ ending at $j$, respectively, and $I'_{x,j}$, $I_{k,j}^i$, and $D_{k,j}^i$ are the analogous events for the flank-insertion, insertion, and deletion states. The dynamic-programming recurrences follow [1]:

$W_0(i,j)$ : forward probability: sum of the probabilities of all paths from the start/"Begin" state to the anchor point $(i,j)$, i.e. through the event $J_j^i$

$W_1(i,j)$ : backward probability: sum of the probabilities of all paths from $(i,j)$ to the end

$X_{i,j}$ : probability of paths whose last operation at profile position $i$ is a match state

$Y_{k,i,j}$ : probability of paths whose last operation is a deletion state; $k \in \{0,1,2\}$ indexes the 0-nt, 1-nt-emitting, and 2-nt-emitting variants

$Z_{k,i,j}$ : probability of paths whose last operation is an insertion state; $k \in \{0,1,2\}$ indexes the codon-emitting, 1-nt-emitting, and 2-nt-emitting variants

Each DP table at $(i,j)$ is for a particular state $S$ at profile position $i$ (e.g. $X$ is for the match state $M^i$): in the forward pass it represents the total probability of all paths from the start that end in $S$ having emitted the DNA prefix $s_{0..j}$, while in the backward pass the same table represents the total probability of all paths that start from $S$ and reach the end. All $W_0$, $X$, $Y$, and $Z$ cells and border rows and columns are initialized to zero before the forward sweep; untouched borders remain zero throughout. The definition of $Y$ is slightly changed from [1], as the optimizations that warrant the shift in indexing do not cleanly apply to frameshifted alignment.

For any state event $S_j$, let

$$\gamma(S_j) := \sum P(\text{query seq}, \pi \mid \text{HMM}). \tag{13}$$

where the sum is taken over all full alignments in which the event $S_j$ occurs. In particular, $\gamma(M_j^i)$ sums over all alignments that visit $M^i$ while emitting $s_{j-2}s_{j-1}s_j$, and $\gamma(I'_{x,j})$ sums over alignments emitting through the null insertion state $I'_x$ at $j$. This parallels the set $A_{i,j}$ and score $s(i,j)$ of Main Paper Section 2.3: $s(i,j) = \log[\gamma(J_j^i)/P(\text{query seq} \mid \text{null})]$, but $\gamma$ itself is a raw probability, with no logarithm and no division by the null probability. All $\gamma(S_j)$ below are obtainable from $W_0$ and $W_1$ (with the state probabilities $X$, $Y$, $Z$ supplying the through-state component for emitting states), as computed by the forward and backward recurrences.

The prefix probabilities $P(s_{0..j} \mid \text{null})$ are those of Appendix S2, precomputed in $O(n)$ time before the forward pass.

#### S3.2 Forward recurrences

We write the recurrence for the junction (forward) probability as

$$\begin{aligned} w_k = X_{i-1,\,j-k} &+ \sum_{d=0}^{2} Y_{d,\,i-1,\,j-k} + \sum_{d=0}^{2} Z_{d,\,i,\,j-k} \\ &+ P(s_{0..j-k} \mid \text{null}) \cdot \left(\tfrac{1}{\sqrt[n]{P(\text{seq}|\text{null})}}\right)^{j-k+1}. \end{aligned} \tag{14}$$

The last term represents starting a new alignment at position $(i, j-k)$. Insertions use index $i$ while matches and deletions use $i-1$ because insertions do not advance the profile. While stated here with the $n$th-root decomposition for clarity, the implementation adds the precomputed null_model_prefix[$j-k$] of Appendix S2 without additional multiplications.

**Match recurrences**

$$X_{i,j} \leftarrow (1 - \sum_k \alpha_{k,i} - \sum_k \delta_{k,i} - \tau) \cdot \theta_i(s_{j-2..j}) \cdot \tfrac{1}{\sqrt[n]{P(\text{seq}|\text{null})}}^{3} \cdot w_3. \tag{15}$$

**Insertion recurrences**

$$\begin{aligned} Z_{0,i,j} &\leftarrow \psi(s_{j-2..j}) \cdot \tfrac{1}{\sqrt[n]{P(\text{seq}|\text{null})}}^{3} \Big(\alpha_{0,i}(1-\beta_{0,i})w_3 \\ &\qquad + \beta_{0,i} Z_{0,i,j-3} + \sum_{k=1}^{2} \frac{\beta_{k,i}(1-\beta_{0,i})}{1-\beta_{k,i}} Z_{k,i,j-3}\Big), \\ Z_{1,i,j} &\leftarrow \alpha_{1,i} \cdot \phi(s_j) \cdot \tfrac{1}{\sqrt[n]{P(\text{seq}|\text{null})}}^{1} \cdot (1-\beta_{1,i}) \cdot w_1, \\ Z_{2,i,j} &\leftarrow \alpha_{2,i} \cdot \phi(s_{j-1}) \cdot \phi(s_j) \cdot \tfrac{1}{\sqrt[n]{P(\text{seq}|\text{null})}}^{2} \cdot (1-\beta_{2,i}) \cdot w_2. \end{aligned} \tag{16}$$

**Deletion recurrences**

$$
\begin{aligned}
Y_{0,i,j} &\leftarrow \delta_{0,i}(1-\varepsilon_{0,i+1})w_0 + \sum_{k=0}^{2} \frac{\varepsilon_{k,i}(1-\varepsilon_{k,i+1})}{1-\varepsilon_{k,i}} Y_{k,i-1,j}, \\
Y_{1,i,j} &\leftarrow \delta_{1,i}\cdot\phi(s_{j-1})\cdot\phi(s_j)\cdot \frac{1}{\sqrt[n]{P(\text{seq}|\text{null})}}^{2}(1-\varepsilon_{1,i+1})w_2, \\
Y_{2,i,j} &\leftarrow \delta_{2,i}\cdot\phi(s_j)\cdot \frac{1}{\sqrt[n]{P(\text{seq}|\text{null})}}^{1}(1-\varepsilon_{2,i+1})w_1.
\end{aligned}
\tag{17}
$$

#### S3.3 Backward recurrences

The backward pass computes the probability of all paths from each state $(i, j)$ to the end, producing the backward (suffix) probability $W_1$, with null scaling as in Appendix S2. Below, $X_{i,j}, Y_{k,i,j}, Z_{k,i,j}$ denote the backward counterparts of the forward DP probabilities in Appendix S3.2 (suffix probabilities from each emitting state back to End, rather than prefix probabilities from Begin), computed in place.

The recurrences hold for $0 \leq i \leq m$ and $0 \leq j < n$. All $W$, $X$, $Y$, and $Z$ cells are initialized to zero before the backward sweep. There is an extra profile position $i = m$ in [1], representing the possibility of having insertions after the profile. The suffix probabilities $P(s_{j..n-1} \mid \text{null})$ are those of Appendix S2, precomputed in $O(n)$ time via a reverse pass, entering only through the null_model_suffix array of Appendix S2.

We write the recurrence for the junction (backward) probability as

$$
\begin{aligned}
W_1(i,j) \leftarrow\ & (1-\sum_k \alpha_{k,i} - \sum_k \delta_{k,i} - \tau)\theta_i(s_{j+1..j+3})\cdot \frac{1}{\sqrt[n]{P(\text{seq}|\text{null})}}^{3}\cdot X_{i+1,j+3} \\
& + \delta_{0,i}\cdot(1-\varepsilon_{0,i+1})Y_{0,i+1,j} \\
& + \delta_{1,i}\cdot(1-\varepsilon_{1,i+1})\cdot\phi(s_{j+1})\cdot\phi(s_{j+2})\cdot \frac{1}{\sqrt[n]{P(\text{seq}|\text{null})}}^{2}\cdot Y_{1,i+1,j+2} \\
& + \delta_{2,i}\cdot(1-\varepsilon_{2,i+1})\cdot\phi(s_{j+1})\cdot \frac{1}{\sqrt[n]{P(\text{seq}|\text{null})}}\cdot Y_{2,i+1,j+1} \\
& + \alpha_{0,i}\cdot(1-\beta_{0,i})\cdot\psi(s_{j+1..j+3})\cdot \frac{1}{\sqrt[n]{P(\text{seq}|\text{null})}}^{3}\cdot Z_{0,i,j+3} \\
& + \alpha_{1,i}\cdot(1-\beta_{1,i})\cdot\phi(s_{j+1})\cdot \frac{1}{\sqrt[n]{P(\text{seq}|\text{null})}}\cdot Z_{1,i,j+1} \\
& + \alpha_{2,i}\cdot(1-\beta_{2,i})\cdot\phi(s_{j+1})\cdot\phi(s_{j+2})\cdot \frac{1}{\sqrt[n]{P(\text{seq}|\text{null})}}^{2}\cdot Z_{2,i,j+2} \\
& + P(s_{j+1..n-1} \mid \text{null})\cdot\left(\frac{1}{\sqrt[n]{P(\text{seq}|\text{null})}}\right)^{\text{n-1-j}}.
\end{aligned}
\tag{18}
$$

**Match recurrences**

$$
X_{i,j} \leftarrow W_1(i,j). \tag{19}
$$

**Insertion recurrences**

$$
\begin{aligned}
Z_{0,i,j} &\leftarrow W_1(i,j) + \beta_{0,i}\cdot\psi(s_{j+1..j+3})\cdot \frac{1}{\sqrt[n]{P(\text{seq}|\text{null})}}^{3}\cdot Z_{0,i,j+3}, \\
Z_{1,i,j} &\leftarrow W_1(i,j) + \frac{\beta_{1,i}(1-\beta_{0,i})}{1-\beta_{1,i}}\cdot\phi(s_{j+1})\cdot \frac{1}{\sqrt[n]{P(\text{seq}|\text{null})}}\cdot Z_{0,i,j+1}, \\
Z_{2,i,j} &\leftarrow W_1(i,j) + \frac{\beta_{2,i}(1-\beta_{0,i})}{1-\beta_{2,i}}\cdot\phi(s_{j+1})\cdot\phi(s_{j+2})\cdot \frac{1}{\sqrt[n]{P(\text{seq}|\text{null})}}^{2}\cdot Z_{0,i,j+2}.
\end{aligned}
\tag{20}
$$

**Deletion recurrences**

$$
\begin{aligned}
Y_{0,i,j} &\leftarrow W_1(i,j) + \frac{\varepsilon_{0,i}(1-\varepsilon_{0,i+1})}{1-\varepsilon_{0,i}} Y_{0,i+1,j}, \\
Y_{1,i,j} &\leftarrow W_1(i,j) + \frac{\varepsilon_{1,i}(1-\varepsilon_{1,i+1})}{1-\varepsilon_{1,i}}\cdot Y_{0,i+1,j}, \\
Y_{2,i,j} &\leftarrow W_1(i,j) + \frac{\varepsilon_{2,i}(1-\varepsilon_{2,i+1})}{1-\varepsilon_{2,i}}\cdot Y_{0,i+1,j}.
\end{aligned}
\tag{21}
$$

This pass also computes the Forward-Backward scores $s(i, j)$ defined in the Main Paper (Section 2.3). For all $i, j$, $s(i, j) = \log[W_0(i, j)W_1(i, j)] = \log[\gamma(J^i_j)/P(\text{seq} \mid \text{null})]$.

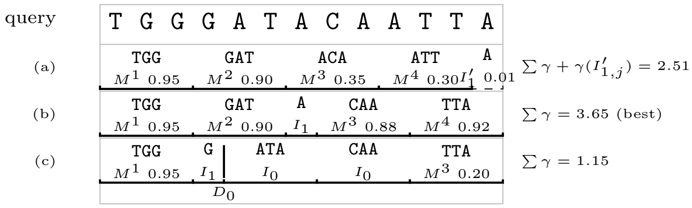


Figure S3: Three valid alignments (figure labels are 1-based for readability; the text is 0-indexed). (a) Forced all-match alignment without frameshifts, giving weak posteriors after a 1-nt insertion; the unaligned terminal base is scored as $\gamma(I'_{1,j})$. (b) One 1-base insert absorbs the extra A, resulting in better succeeding matches. (c) Arbitrary alignment with suboptimal scores. Match labels give the posterior weights $\gamma(M^i_j)$. If these were the set of alignments available, we select alignment (b).

### S4 Representative alignment and range dynamic programming

The representative alignment is a path $\pi^*$, a local alignment lying in the open interval $(l, r)$ of the query sequence, where the alignment enters from the null model after position $l$ and exits at position $r$ (the flanks are $j \le l$ and $j \ge r$). We find the $\pi^*$ maximizing

$$\sum_{M^i_j \in \pi^*} 3\,\gamma(M^i_j) \;+\; \sum_{x\in\{1,2,3\}} x\Big(\sum_{j\le l}\gamma(I'_{x,j}) + \sum_{j\ge r}\gamma(I'_{x,j})\Big), \tag{22}$$

where $\gamma(I'_{x,j})$ is the posterior probability of the null insertion state $I'_x$ (emitting $x$ nucleotides, $1 \le x \le 3$) at positions outside $(l, r)$; each posterior probability is weighted by the number of nucleotides it emits. Insertions and deletions within the alignment are chosen implicitly as the connections between selected matches.

We denote the two flank sums by $L_l$ and $R_r$ ($L_j := \sum_{k\le j}\sum_x x\,\gamma(I'_{x,k})$ and $R_j := \sum_{k\ge j}\sum_x x\,\gamma(I'_{x,k})$). The objective can then be re-expressed as

$$\sum_{M^i_j \in \pi^*} 3\,\gamma(M^i_j) \;+\; L_l \;+\; R_r. \tag{23}$$

#### S4.1 Dynamic Programming recurrences

The prefix-optimal matrix $\Gamma^{\text{pre}}$ is built with:

$$\Gamma^{\text{pre}}(i,j) = \max\{\Gamma^{\text{pre}}(i-1,j),\ \Gamma^{\text{pre}}(i,j-1),\ 3\gamma(M^i_j) + \Gamma^{\text{pre}}(i-1,j-3),\ L_j\}, \tag{24}$$

with options deletion, insertion, match emission, and null entry with $l = j$ ($L_j$ as defined above). The implementation evaluates $L_j$ and $R_j$ by column-wise probability accumulation with frameshift spreading and opposite-direction cumulative sums, in null-absorbed scaled units; the uniform scale leaves the arg max unchanged. The suffix-optimal matrix $\Gamma^{\text{suf}}$ is built with:

$$\Gamma^{\text{suf}}(i,j) = \max\{\Gamma^{\text{suf}}(i+1,j),\ \Gamma^{\text{suf}}(i,j+1),\ 3\gamma(M^i_j) + \Gamma^{\text{suf}}(i+1,j+3),\ R_j\}, \tag{25}$$

with options deletion, insertion, match emission, and null exit with $r = j$.

Alignments are obtained by performing tracebacks on these dynamic programming matrices.
